# KCF-Convoy: efficient Python package to convert KEGG Chemical Function and Substructure fingerprints

**DOI:** 10.1101/452383

**Authors:** Masayuki Sato, Hirotaka Suetake, Masaaki Kotera

## Abstract

**Motivation:** *In silico* methodologies to assess pharmaceutical activity and toxicity are increasingly important in QSAR, and many chemical fingerprints have been developed to tackle this problem. Among them, KEGG Chemical Function and Substructure (KCF-S) has been shown to perform well in some pharmaceutical and metabolic studies. However, the software that generates KCF-S fingerprints has limited usability: the input file must be Molfile or SDF format, and the output is only a text file.

**Results:** We established a new Python package, KCF-Convoy, to generate KCF format and KCF-S fingerprints from Molfile, SDF, SMILES, and InChI seamlessly. The obtained KCF-S was used in a number of supervised machine-learning methods to distinguish herbicides from other pesticides, and to find characteristic substructures in taxonomy groups.

**Availability:** KCF-Convoy is implemented as a Python package freely available at https://github.com/KCF-Convoy and the user can use the package management system “pip” and also the Docker environment.

## 1 Introduction

Functionality and toxicity are inseparable characteristics of chemical compounds, necessitating the effective estimation of safety for the development of pharmaceuticals and bioactive compounds (Roy et al., 2015). To reduce the research and development costs of such compounds, and also from the viewpoint of animal welfare, there is an increasing need to develop *in vitro* and *in silico* methodologies that can replace conventional toxicity tests. To this end, it is considered beneficial to use animal experiment data that are already available, and to apply artificial intelligence and machine learning techniques to obtain brand new knowledge on toxicology. These approaches should help to achieve high-accuracy toxicity prediction methods covering various types of compounds that cannot be studied using existing Quantitative Structure-Activity Relationship (QSAR) models.

For computational analyses, chemical structures can be represented in various formats; e.g., Molfile, SDF, SMILES, InChI (Heller et al., 2015), and KEGG Chemical Function (KCF) (Hattori et al., 2003) that can be irreversibly transformed into numeric vectors, sometimes referred to as chemical fingerprints. They are used in a wide range of studies, including chemical similarity searches, QSAR for drug discovery, and metabolic pathway analyses. Many fingerprints are available, including MACCS (Durant et al., 2002), Pub Chem (Chen et al., 2009), and KCF and Substructure (KCF-S) (Kotera et al., 2013). Many chemical toolkits, e.g., OpenBabel, RDKit, CDK, Daylight, OpenEye, Indigo, and Frowns, are available to calculate various types of chemical fingerprints.

Among them, KCF-S has been shown to perform the best in some pharmaceutical and metabolic studies (Kotera et al., 2013; Sawada et al., 2014). However, there are currently only two ways to generate KCF format, namely via the chemical analysis tools on the GenomeNet API (http://www.genome.jp/tools/gn_ca_tools_api.html), and the downloadable KCFCO software provided by GenomeNet (http://www.genome.jp/en/gn_ftp.html). Both these ways have limited usability: the input file must be only Molfile or SDF format, and the output is only a text file.

Here, we established a new Python package, named KCF-Convoy, to generate KCF format and KCF-S fingerprints from Molfile, SDF, SMILES, and InChI files seamlessly (Figure 1). The procedure developed by Kotera et al., 2013 (Figure 1a) requires several steps that generate intermediary files, and the resulting KCF-S fingerprints are obtained as a text file. KCF-Convoy (Figure 1b) can generate KCF-S fingerprints without producing intermediary text files. The KCF-S fingerprints are generated in NumPy arrays (http://www.numpy.org/), and can be directly subjected to the scikit-learn machine-learning library (http://scikit-learn.org/) and other Python libraries. We also added options to select KCF-S attributes that represent different levels of chemical substructures (Kotera et al., 2013). The KCF-Convoy package should improve the usability of KCF-S fingerprints and further contribute to chemical bioinformatics studies.

**Fig. 1.**
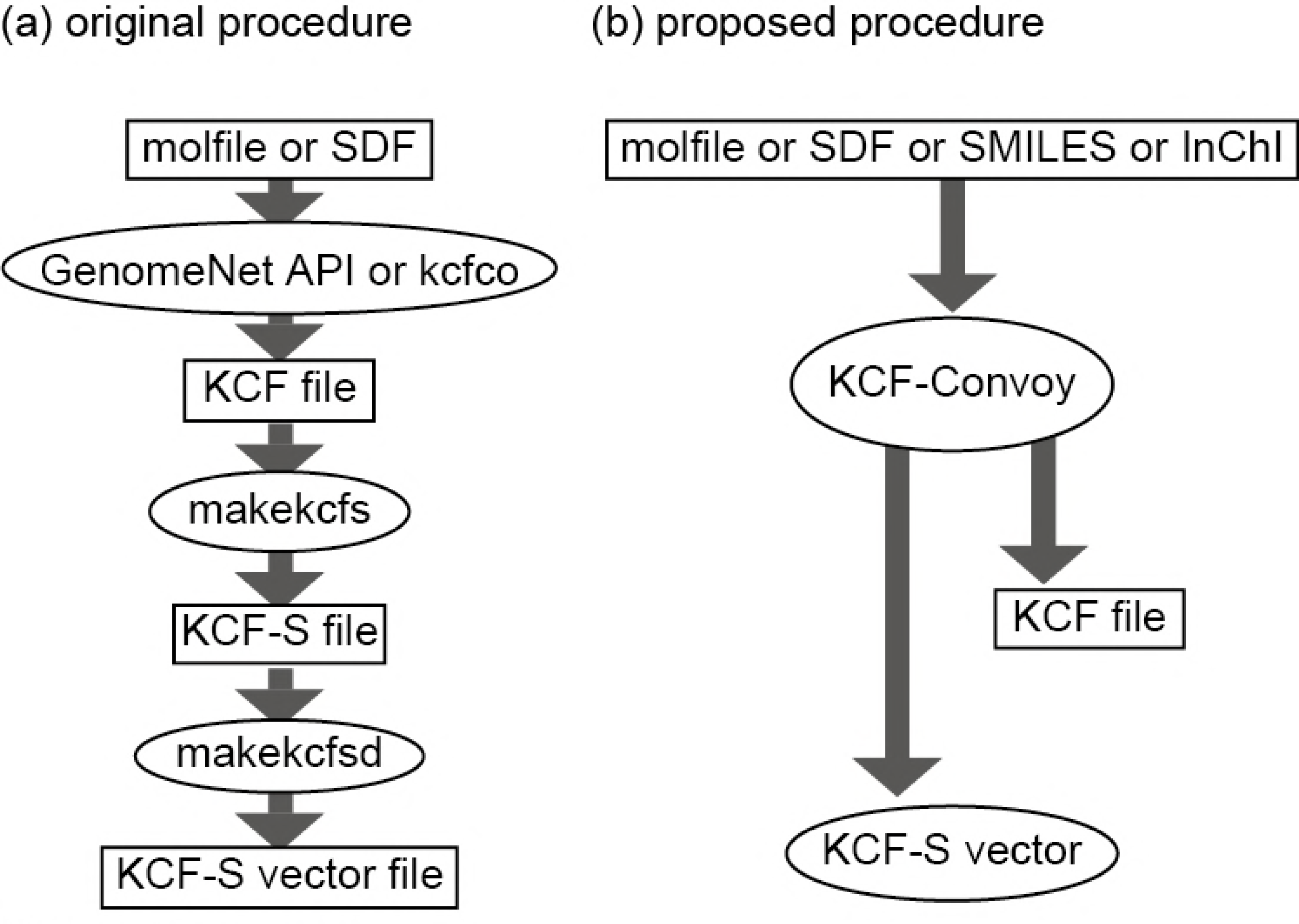
Differences between (a) the original and (b) proposed procedures for generating KCF Substructure (KCF-S) vectors. Ellipses and rectangles indicate executable programs and text files, respectively.

## 2 Design and implementation

We prepared a variety of installation methods as explained in the Wiki page of https://github.com/KCF-Convoy/. One option is to use pip, the Python package management system. The RDKit library should be installed in advance. Miniconda (https://conda.io/miniconda.html) is recommended for this purpose. The following pip command can then be used to install the KCF-Convoy package:

% pip install kcfconvoy

It is also recommended to use the Docker environment to install KCF-Convoy as explained in the KCF-Convoy wiki.

In the KCF-Convoy package, the users should handle the two modules:

1. kcfconvoy.KCFvec - defines a KCF-S fingerprint (a compound)
2. kcfconvoy.KCFmat - defines a group of KCF-S fingerprints

KCF-Convoy uses the RDKit (http://www.rdkit.org/) and NetworkX (https://networkx.github.io/) packages. Different from the original KCF Converter in GenomeNet, KCF-Convoy enables molecules to be input not only as Molfile and SDF files, but also as SMILES and InChI files. It also enables the conversion of KCF format to other formats such as Molfile.

KCF-Convoy depicts a molecule as a mathematical graph structure and calculates KCF-defined biochemical substructures (Kotera et al., 2013), where vertices (nodes) are assigned KEGG atom labels that represent atomic elements with hybrid orbit and other physicochemical information (Hattori et al., 2003). KEGG atom labels generally consist of three letters; for example, C1a, where C is the element symbol of carbon, C1 indicates a sp3 carbon, and C1a indicates a methyl carbon (an sp3 carbon that forms a bond with only one atom other than hydrogen atoms). Definitions of all KEGG atoms are listed in http://www.genome.jp/kegg/reaction/KCF.html. KCF-Convoy provides an integer KCF vector if a query molecule is given, and an integer KCF matrix if a group of molecules are given.

## 3 Applications

It has been shown previously that KCF-S vectors can outperform other chemical fingerprints in drug–target interaction prediction (Sawada et al., 2014) and enzymatic reaction prediction (Kotera et al., 2013). In this paper, we demonstrate two other examples to show the usefulness of KCF-S vectors.

### 3.1 Supervised machine learning to distinguish herbicides from other pesticides

An advantage of KCF-S compared to other chemical fingerprints is its interpretability. Figure 2 shows examples of molecules and their KCF Substructures that were found specifically either in herbicides or in other pesticides registered in the KEGG database (The complete list of 918 pesticides was obtained from http://www.kegg.jp/keggbin/get_htext?br08007.keg). The C1a-O2a-C8y-N5x-C8y substructure was found only in herbicides, whereas the S0-P1a-O2b-C1b-C1a substructure was found in insecticides and fungicides, but not in herbicides. These examples show that KCF-S enables the interpretation of substructures that contribute to the molecular functions of the corresponding molecules.

**Fig. 2.**
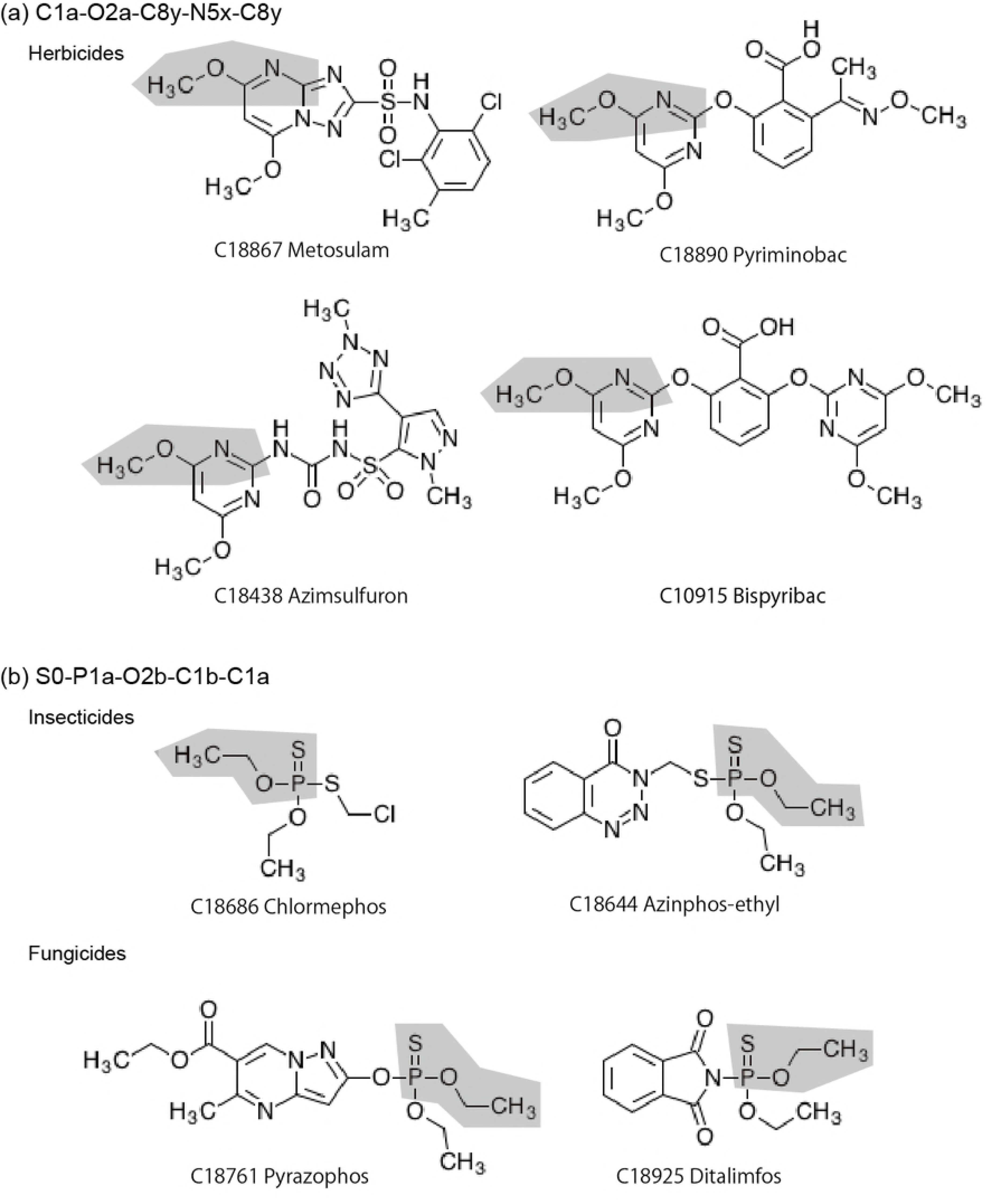
Examples of substructures found specifically in (a) herbicides and (b) other pesticides. The C1a-O2aC8y-N5x-C8y and S0-P1a-O2b-C1b-C1a strings describe substructures in KCF-S, and are highlighted in gray.

To distinguish herbicides from other pesticides, we used different supervised machine-learning methods in the scikit-learn package, namely logistic regression, kNN (k-nearest neighbors), linear SVM (support vector machine), polynomial SVM, RBF (radial basis function) SVM, sigmoid SVM, decision tree, random forest, AdaBoost, extra tree, Gaussian process, gradient boosting, SGD (stochastic gradient decent), LDA (linear discriminant analysis), QDA (quadratic discriminant analysis), MLP (multi-layer perceptron) and naive Bayes. We used the KEGG BRITE database (Kanehisa et al., 2016) to collect the chemical structures of 918 pesticides, which included 365 herbicides, 299 insecticides, and 240 fungicides.

Table 1 shows the summary of the results for the supervised machine-learning methods using KCF-S, Morgan, RDKit, Layered, and Pattern fingerprints as inputs. For most machine-learning methods, KCF-S generated better prediction results than the other four fingerprints. See https://github.com/KCF-Convoy/kcfconvoy/wiki/Example-usage-of-KCF-Convoy-for-machine-learning for the detailed results and the Python code.

**Table 1.** Accuracy scores for the supervised machine-learning methods and chemical descriptors used to distinguish herbicides from other pesticides. The best prediction for each machine-learning methods are shown in bold letters.

| Fingerprint | KCF-S | Morgan | RDk | Layered | Pattern |
| --- | --- | --- | --- | --- | --- |
| Dimension | 2,048 | 2,048 | 2,048 | 2,048 | 1,024 |
| MLP | **0.859 ± 0.017** | 0.833 ± 0.014 | 0.833 ± 0.019 | 0.814 ± 0.016 | 0.810 ± 0.018 |
| Extra tree | **0.849 ± 0.023** | 0.815 ± 0.014 | 0.813 ± 0.011 | 0.811 ± 0.013 | 0.811 ± 0.011 |
| Logistic regression | **0.840 ± 0.019** | 0.835 ± 0.012 | 0.831 ± 0.018 | 0.813 ± 0.023 | 0.786 ± 0.017 |
| Gradient boosting | **0.834 ± 0.013** | 0.800 ± 0.019 | 0.827 ± 0.015 | 0.807 ± 0.013 | 0.805 ± 0.017 |
| Gaussian process | 0.831 ± 0.016 | **0.844 ± 0.012** | 0.838 ± 0.013 | 0.826 ± 0.022 | 0.794 ± 0.011 |
| Random forest | **0.827 ± 0.017** | 0.806 ± 0.015 | 0.791 ± 0.015 | 0.795 ± 0.018 | 0.798 ± 0.021 |
| Linear SVM | 0.816 ± 0.018 | **0.834 ± 0.015** | 0.804 ± 0.014 | 0.785 ± 0.016 | 0.766 ± 0.023 |
| AdaBoost | **0.805 ± 0.016** | 0.770 ± 0.012 | 0.759 ± 0.017 | 0.782 ± 0.019 | 0.754 ± 0.017 |
| kNN | 0.794 ± 0.017 | **0.796 ± 0.019** | 0.782 ± 0.028 | 0.789 ± 0.016 | 0.767 ± 0.019 |
| SGD | 0.792 ± 0.017 | 0.800 ± 0.021 | **0.801 ± 0.020** | 0.787 ± 0.021 | 0.714 ± 0.072 |
| Naïve Bayes | **0.789 ± 0.014** | 0.756 ± 0.024 | 0.745 ± 0.035 | 0.672 ± 0.038 | 0.526 ± 0.021 |
| Decision tree | **0.785 ± 0.012** | 0.772 ± 0.018 | 0.729 ± 0.020 | 0.743 ± 0.035 | 0.742 ± 0.027 |
| LDA | **0.779 ± 0.024** | 0.697 ± 0.046 | 0.759 ± 0.029 | 0.744 ± 0.024 | 0.644 ± 0.023 |
| RBF SVM | **0.770 ± 0.017** | 0.639 ± 0.017 | 0.761 ± 0.017 | 0.690 ± 0.034 | 0.679 ± 0.024 |
| QDA | **0.748 ± 0.050** | 0.557 ± 0.100 | 0.637 ± 0.058 | 0.581 ± 0.065 | 0.582 ± 0.139 |
| Sigmoid SVM | 0.666 ± 0.020 | 0.639 ± 0.017 | **0.694 ± 0.030** | 0.667 ± 0.024 | 0.651 ± 0.026 |
| Polynomial SVM | 0.637 ± 0.012 | 0.639 ± 0.017 | **0.645 ± 0.019** | 0.638 ± 0.022 | 0.649 ± 0.022 |

### 3.2 Extraction of characteristic substructures in a genus

Based on the assumption that taxonomically related species conserve similar metabolic pathways and chemical substructures, we investigated the appearance of substructures in the metabolite–species relationships described in the KNApSAcK database (Nakamura et al., 2014). As a result, we detected chemical substructures that were specific in a particular genus. We found that the N1b-C2c-S2a substructure was present only in eight metabolites out of the 50899 metabolites in KNApSAcK (Figure 3), and four of them were specifically present in genus *Brassica*.

**Fig. 3.**
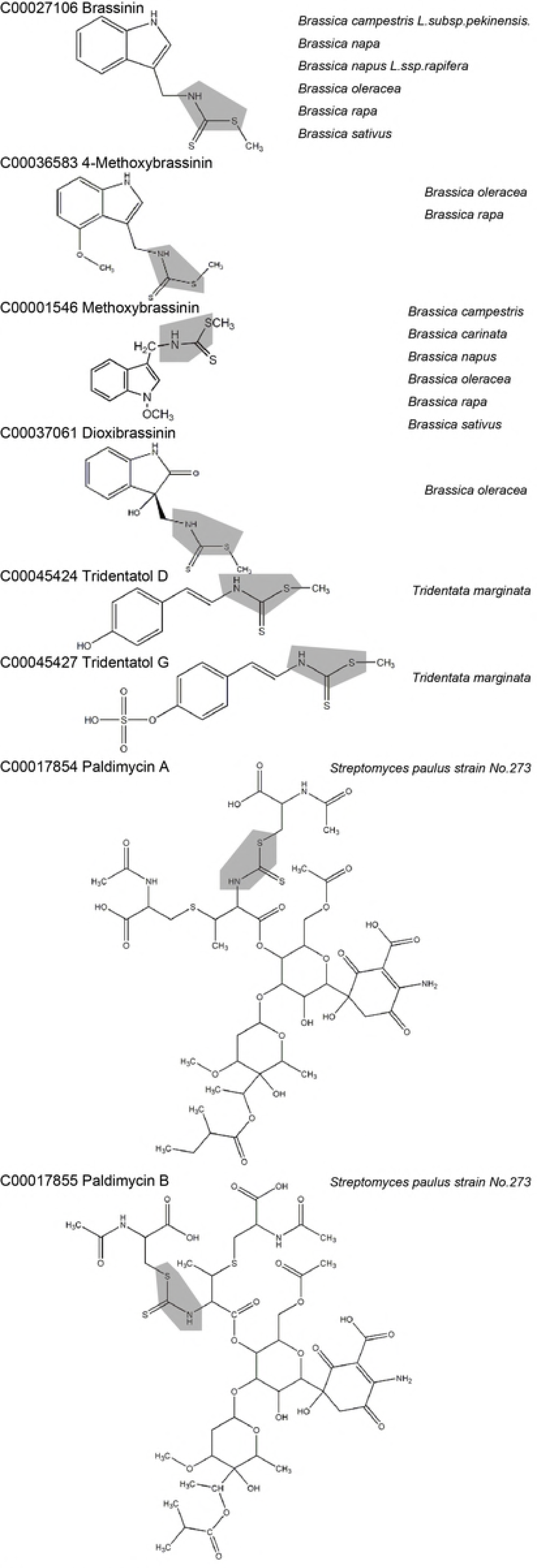
Metabolites that contain a N1b-C2c-S2a substructure. The N1b-C2c-S2a substructures are highlighted in gray. Species that possess the metabolites are in italics.

The eight metabolites were subjected to the E-zyme2 program (Moriya et al., 2016), which predicts candidate enzyme orthologs that can catalyze a reaction given as a query substrate–product pair. The prediction results showed that a *Brassica oleracea* (wild cabbage) cytochrome (CYP) enzyme (EC 1.14.14.1) could catalyze the interconversion of two of the metabolites (C00027106 and C00036583). We considered this reasonable because the difference between these two metabolites is one methoxy group (-OCH_3_), indicating the hydroxylation reaction (typically catalyzed by CYP) may be followed by the methylation reaction.

The substructure effects in ligand-protein interactions have been investigated systematically and comprehensively (e.g., Malhotra and Karanicolas, 2017; Korkuc and Walther, 2016). This study reinforces the potential of KCF-S for *de novo* pathway reconstruction of natural products, and should also allow further understanding of substructure effects in wider range of issues in molecular biology.

## Acknowledgment

We thank Margaret Biswas, PhD, from Edanz Group (www.edanzediting.com/ac) for editing a draft of this manuscript.

